# ACAN regulates VSMC mitochondrial homeostasis through the YAP1/TAZ pathway and contributes to the onset and progression of aortic dissection

**DOI:** 10.64898/2026.09.14.751622

**Authors:** Wenjie Fu, Yanlan Gao, Yanqing Fang, Linlong Guo, Yalu Huang, Luyao Hong, Binxing Lu, Ziru Li, Laixiang Yan, Bin Hu, Chaoyun Wang, Lin Zhong, Min Chen, Yumei Li

## Abstract

**Objective:** Aortic dissection (AD) is a serious, life-threatening cardiovascular crisis. While vascular proteoglycans normally buffer hemodynamic stress, their aberrant accumulation in the AD aorta predisposes to rupture. Aggrecan (ACAN) is among the most frequently elevated proteoglycans in this setting, yet its pathogenic relationship to AD remains undefined. This study aimed to investigate the role of ACAN in AD and identify potential therapeutic targets.

**Approach and Results:** We employed human tissues, animal models, and cell cultures. Western blotting and immunohistochemistry (IHC) were performed on aortic samples from AD patients and controls to assess contractile markers, MMPs, ACAN, and SOX9. In vivo, AD was induced in mice using β-aminoisobutyric acid (BAPN) and Ang II, with AAV-mediated ACAN knockdown in VSMCs. Histological staining confirmed model validity and enabled assessment of AD incidence, mortality, and protein expression changes. In vitro, human aortic vascular smooth muscle cells (HAVSMCs) were stimulated with Ang II, and SOX9 and ACAN were silenced via siRNA to evaluate the functional impact of ACAN downregulation. In human AD aortas, elevated ACAN and its upstream transcription factor SOX9 drove VSMC phenotypic switching and ECM degradation. In mice, ACAN knockdown corrected AD-induced medial structural disruption, elastin fragmentation, and collagen deposition, markedly reducing incidence and mortality while curbing excessive mitochondrial fission and rescuing functional integrity. ACAN silencing attenuated Ang II-induced mitochondrial damage in HAVSMCs via the RNA-seq-identified YAP1/TAZ pathway, and Verteporfin restored homeostasis.

**Conclusion:** ACAN is aberrantly expressed in AD and murine aortas. ACAN downregulation delays disease progression by preserving mitochondrial homeostasis and function via the YAP1/TAZ pathway.

## Introduction

Aortic dissection (AD) has an insidious onset and is life-threatening. It is usually triggered by damage to the aortic intima or rupture of a feeding vessel. This causes blood to rush into the aortic media, tearing the vessel and forming true and false lumens. AD is one of the most dangerous cardiovascular diseases and poses a serious threat to the health and lives of the Chinese population. This results in enormous economic losses and places a heavy burden on the healthcare system^1,2^. Following the onset of AD, the mortality rate among patients who do not receive timely clinical intervention rises sharply over time. Compared to medical treatment, surgical intervention remains the primary clinical approach for treating AD. However, due to the lack of specific clinical signs and the limited sensitivity of routine imaging tests such as ECGs and CT scans, most AD patients struggle to receive timely treatment ^3–5^. Therefore, investigating the pathophysiological mechanisms of AD, identifying novel diagnostic biomarkers, and discovering potential drug targets are current priorities in AD research.

The pathogenesis of AD is complex, and mid-aortic wall pathology is considered one of the primary causes of AD. The mid-aortic wall is composed of vascular smooth muscle cells (VSMCs) and the extracellular matrix (ECM), which together maintain vascular mechanical function and help the aorta resist shear stress from blood flow ^6,7^. The ECM plays a crucial role in maintaining the structural and functional integrity of the aorta; it regulates VSMC function and activity through various signalling pathways. Once ECM homeostasis is disrupted, it may trigger AD development. Most currently identified genetic factors, such as Marfan syndrome and Loeys-Dietz syndrome, are associated with ECM alterations that significantly increase the risk of AD ^8–10^.

Proteoglycans are components of the ECM. These complex molecules are composed of a core protein and various covalently linked glycosaminoglycan (GAG) chains. They exhibit a high negative charge and strong hydrophilicity. By intertwining with collagen, they form a compression-resistant extracellular matrix. Through electrostatic attraction, they draw in active ions to maintain vascular elasticity. This plays a crucial role in preserving the structural and functional stability of arteries ^11^. While the role of proteoglycans in AD has often been overlooked in past studies, recent findings suggest that abnormal accumulation of proteoglycans in the aortic wall may contribute to AD ^12,13^. Mass spectrometry identified aggrecan (ACAN) and versican (VCAN) as the predominant proteoglycans markedly elevated in the aortas of patients with AD ^14^. Subsequent studies have found that VCAN promotes AD development by inducing aortic wall dilation in Marfan syndrome mice through activation of the Akt pathway ^15^. Furthermore, multiple studies have reported on the role of the ADAMTS family, which degrades ACAN and VCAN in blood vessels, in the onset and progression of AD ^16,17^. However, further research is needed to elucidate the role of ACAN in AD.

VSMCs are the primary component of the aortic media, and those with a contractile phenotype play a crucial role in maintaining aortic elasticity and withstanding blood flow stress ^18^. Pathological changes in VSMCs are strongly associated with cardiovascular diseases, including atherosclerosis and pulmonary arterial hypertension^19^. Studies on the aortas of patients with AD have revealed a significant loss of VSMCs in the aortic wall. VSMCs with a contractile phenotype, which are responsible for maintaining vascular function, are converted to a synthetic phenotype and secrete increased levels of matrix metalloproteinases (MMPs), which further exacerbate ECM remodelling ^20,21^. These studies confirm that VSMC dysfunction is a core pathological mechanism in the development and progression of AD. Understanding the mechanisms underlying VSMC changes during the onset and progression of AD could help to develop early prevention strategies and novel targeted therapies, thereby improving the current treatment landscape for AD. Mitochondria are the centres of cellular energy metabolism and play a crucial role in maintaining the normal contractile phenotype and physiological activities of VSMCs. Mitochondrial dysfunction and oxidative stress lead to the loss of VSMCs and accelerate degenerative changes in the vascular wall ^22^. A key factor in the pathogenesis of cardiovascular disease is the transition of mitochondrial morphology from normal to fragmented or abnormally dilated, particularly an imbalance between excessive fission and fusion ^23^. Normal mitochondria perform multiple vital functions, including the production of ATP, the maintenance of calcium homeostasis, and the metabolism of amino acids and fatty acids. However, an imbalance in mitochondrial dynamics can significantly impair ATP production and disrupt redox balance and intracellular calcium homeostasis. This can exacerbate oxidative stress and inflammatory responses, inducing cell necrosis and apoptosis ^24^. In our previous study, we found that before and after the onset of AD, homeostasis of mitochondrial fission and fusion within VSMCs is disrupted, mitochondrial fission is exacerbated, and expression of proteins associated with fusion of the inner and outer mitochondrial membranes, such as MFN1 and OPA1, is suppressed. This ultimately leads to loss of normal mitochondrial morphology and function, contributing to loss of AD ^25^. Furthermore, recent studies have shown that ECM damage caused by infection or mechanical stress may trigger mitochondrial-mediated adaptive immune and metabolic responses, suggesting a potential association between ECM remodelling and mitochondrial homeostasis ^26,27^.

Taken together, the pathogenesis of AD is highly complex. Current research suggests that VSMC phenotypic switching, programmed cell death, and ECM remodelling may not be isolated events but are closely interrelated.

In this study, we aim to discuss the mechanism of action of ACAN in AD. In our initial research, we confirmed elevated ACAN expression in the aortic media of AD patients, suggesting that ACAN may be involved in regulating AD progression. Furthermore, studies using a BAPN/Ang II-induced AD mouse model and an Ang II-induced in vitro VSMC AD model revealed that ACAN may induce AD by disrupting mitochondrial homeostasis. Finally, using transcriptomics and adenoviral intervention, we revealed that excessive ACAN accumulation may promote mitochondrial dysregulation via the YAP1/TAZ pathway, thereby mediating the onset of AD. These findings are expected to provide new insights into the pathological mechanisms of AD and into drug development.

## Materials and methods

### Human specimens

Aortic tissue from patients with AD and the control group was provided by Union Hospital, affiliated with Fujian Medical University. This included five cases in the control group and five in the AD patient group. The control group consisted of normal thoracic aortic tissue from heart transplant donors with no history of cardiovascular disease. Samples from the AD patient group were obtained from patients with AD who underwent artificial vascular replacement surgery. All patients underwent a review of their clinical history, a physical examination, and an imaging diagnosis. Patient information is summarised in Table 1. This research project was approved by the Ethics Committee of Fujian Medical University and Xiehe Hospital of Fujian Medical University (No. 2024-42) and conducted under their guidance. All participants signed written informed consent forms.

### Animal experiments

Healthy, three-week-old, male C57BL/6J mice were supplied by Shanghai Slaike Laboratory Animal Co., Ltd. Laboratory Animal Production License Number: SCXK (Hu) 2022-0004. All animal experimental protocols complied with the Guidelines for the Care and Use of Laboratory Animals in China, and this study was approved by the Institutional Animal Care and Use Committee (IACUC) of Fujian Medical University (IACUC FJMU 2024-Y-0367). Mice were fed 0.3% BAPN in their drinking water for four weeks. Starting on day 36, Ang II was administered via subcutaneous injection at a total dose of 1.44 mg/kg twice daily at six-hour intervals. Three days later, the surviving mice were euthanized with an overdose of urethane, and their aortas were harvested. Following histological staining, the formation of false lumens in the aortic wall was examined under a microscope.

### Pretreatment of human aortic wall tissue

Patients with AD were selected after excluding conditions such as Marfan syndrome, bicuspid aortic valve, syphilitic aneurysm, and arteritis. Aortic samples in the control group were obtained from heart transplant donors without cardiovascular disease. Following removal, the patients’ aortic tissue was immersed in saline and transported to the laboratory within 30 minutes. A portion of the tissue was blotted dry and stored at −80 °C for subsequent experiments. The remaining tissue was trimmed into several 1 cm strips and fixed in 4% paraformaldehyde to facilitate subsequent procedures, such as tissue dehydration, embedding, and histological staining.

### Construction and applications of adeno-associated virus (AAV)

The shACAN gene from the mouse was cloned into an AAV9 plasmid and packaged into an AAV9 virus. This virus was then used to transfect mouse aortic VSMCs, reducing ACAN expression in these cells. The shACAN and control adeno-associated viral vectors were designed by Shanghai Hanheng Biotechnology Co., Ltd. in accordance with the protocol. The plasmid was transfected into AAV-293 cells with Lipofiter™. Seventy-two hours later, cells with AAV particles were scraped, centrifuged, and the lysate supernatant containing AAV was collected. After treatment with All-Purpose Nuclease and purification, the viral sample was ultrafiltered and collected. The virus was tested for sterility, mycoplasma, and titer; results were: HBAAV2/9-sm22a-ZsGreen control: 1.9×10¹² vg/mL; HBAAV2/9-sm22a-Mir30-mAcan-ZsGreen: 2.0×10¹² vg/mL. AAV9 was injected via the tail vein one week before the mice consumed 0.3% BAPN water, and infection efficiency was assessed after three weeks.

### Cell culture

HAVSMCs were purchased from Shanghai Fuheng Co., Ltd. The cells were cultured in 4 mL of DMEM medium with 1% antibiotics (penicillin and streptomycin) and 10% fetal bovine serum. Incubate at 37 °C. When the cell density reaches 80–90%, pass the cells. Aspirate medium, wash twice with PBS, add 1 mL of 0.25% trypsin (EDTA-free), and observe until cells become round and bright. Add 1 mL of culture medium to stop digestion, then pipette thoroughly. Transfer the suspension to two or three culture flasks, fill each flask with medium as needed, label, and incubate. When density reaches 60-70%, starve in serum-free medium for 24 hours, then switch to medium with 10 μM Ang II for another 24 hours.

### siRNA-mediated gene silencing technology

Cell culture (using a six-well plate as an example; other culture plates or dishes can be adapted accordingly): One day (18-24 hours) before transfection, seed approximately 200,000-700,000 cells per well in a six-well plate. The exact number depends on the cell type, size, and growth rate. This will yield a cell density of approximately 70–80% by the following day. Before proceeding with the transfection steps below, replace the medium in each well of the six-well plate with 1.5 mL of serum-free, antibiotic-free culture medium to improve RNAiMAX transfection efficiency. For each well, take a sterile centrifuge tube and add 250 μL of antibiotic- and serum-free DMEM (high- or low-glucose). Add 100 pmol of siRNA (see Table 5) and mix gently. Take another sterile tube (Tube B), add 250 μL of DMEM, then 5 μL of Lipo3000, and mix gently. Combine tubes A and B, incubate at room temperature for 20 minutes without vortexing or centrifugation. Transfer 500 μL of the mixture to the six-well plate, distribute evenly, and incubate at 37 °C for 6 hours. Remove the mixture, replace it with medium supplemented with 10% FBS, and culture for 48 hours before further experiments.

### Western blot

Total protein from HASMC, the mouse sample, or the human sample was lysed in RIPA buffer as described in detail [25]. Proteins were separated by SDS-PAGE and transferred to PVDF membranes. The membranes were blocked with skimmed milk for 1 h at room temperature and incubated with primary antibodies at 4 °C overnight. After three TBST washes, the membranes were incubated with secondary antibodies for 1 h. The signal was visualized with ECL reagents (36208ES76, Yeason, Shanghai, China), detected with a GE AI 680, and band density quantified using ImageJ. Antibodies used in this study included TAZ (23306-1-AP, Proteintech, Wuhan, China), SM22α (60213-1-lg, Proteintech, Wuhan, China), CNN1 (25905-1-AP, Proteintech, Wuhan, China), β-actin (BA2305, Boster, Wuhan, China), Drp1 (26312-1-AP, Proteintech, Wuhan, China), OPA1 (27733-1-AP, Proteintech, Wuhan, China), YAP1 (81090-1-RR, Proteintech, Wuhan, China), MMP9 (10375-2-AP, Proteintech, Wuhan, China), MMP2 (10373-2-AP, Proteintech, Wuhan, China), Bcl2 (P23612, ProMab Biotechnologies, Hunan, China), Bax (31249, ProMab Biotechnologies, Hunan, China), Sox9 (AF5163, CST, USA), Mfn1 (AF7461, Beyotime, Beijing, China), ACAN (TD7561, Abmart, Shanghai, China), HRP-labeled Goat Anti-Mouse IgG (A0350, Beyotime, Beijing, China), HRP-labeled Goat Anti-Rabbit IgG (A0352, Beyotime, Beijing, China).

### JC-1 Staining for Mitochondrial Membrane Potential

Cultured cells were washed with PBS and incubated with 1 mL of JC-1 staining working solution (C2006, Beyotime) per well at 37 °C for 20 minutes. After incubation, collect the supernatant and wash the cells twice with 1×JC-1 staining buffer. Then, 2 mL of fresh cell culture medium was added, and the mitochondrial membrane potential was evaluated by fluorescence microscope based on JC-1 fluorescence displacement.

### Transmission Electron Microscopy

The sample was fixed with 2.5% glutaraldehyde, and then fixed with 1% yttrium tetraoxide in Millonig phosphate buffer (pH 7.3) for 1 hour. After washing with the same buffer, dehydrate with a graded acetone series. Then the sample is infiltrated, embedded and polymerized with resin. The ultra-thin slice (50-100 nm) was cut on the ultra-micro slice with a diamond knife, double-stained with 3% uranyl acetate and lead citrate, and imaged by transmission electron microscopy.

### RNA-seq

Total RNA was extracted with TRIzol, quantified using a NanoDrop 2000, and its purity assessed. Integrity was assessed with the Agilent 2100 Bioanalyzer. Libraries were prepared with the VAHTS Universal V10 RNA-seq Kit and sequenced on Illumina NovaSeq 6000 for 150 bp paired-end reads. FastP filtered low-quality reads to produce clean data. HISAT2 aligned reads to the reference genome and calculated FPKM. HTseq-count obtained gene counts. PCA analysis and plotting were performed in R to assess replicate quality. DEGs were identified using DESeq2 with q < 0.05 and fold change > 2. Hierarchical clustering and radar charts for the top 30 genes were generated in R using the ggradar package. GO and KEGG enrichment analyses used the hypergeometric distribution. Bar charts, chord diagrams, and pie charts were plotted in R. GSEA software was used to perform gene set enrichment analysis, testing whether predefined gene sets were enriched at either end of the sorted gene list.

### Movat staining

Perform dewaxing of paraffin sections by heating and soaking in dewaxing solution, followed by ethanol washes. Heat Russell’s mordant in a microwave, then treat sections for 10 minutes, rinse for 10 minutes. Add Haibo solution, treat for 5 minutes, rinse with distilled water. Add Alizarin Blue; stain for 20-60 minutes; rinse. Preheat alkaline alcohol to 45-60 °C, incubate the slides for 5-10 minutes, then rinse. Stain with hematoxylin in the dark for 5-10 minutes, differentiate for 20-60 seconds, rinse. Stain with fuchsin for 1 minute, rinse, treat with phosphotungstic acid for 1-2 minutes, then with weak acid for 1-2 minutes. Stain with saffron for 3 minutes, rinse, decolorize, air-dry, clear with xylene, and mount.

### Data analysis

All graphs and calculations were produced using GraphPad Prism 8.0.1 (San Diego, California). Data are presented as the mean ± standard error. The Shapiro-Wilk test was used to assess the normality of the data. When the data were normally distributed, differences between two groups were compared using a two-tailed unpaired t-test; otherwise, the non-parametric Mann-Whitney test was used. Differences among three or more groups were analyzed using one-way ANOVA. If the data were assumed to be normally distributed, Dunnett’s multiple comparison test was used to compare the groups with the control group. Otherwise, the non-parametric Kruskal–Wallis test was used for further analysis. A p-value of less than 0.05 indicates a statistically significant difference (in the figure, p < 0.05 is indicated by *, p < 0.01 by **, and p < 0.001 by ***).

## Result

### The phenotypic transition of aortic VSMCs in patients with AD is accompanied by proteoglycan accumulation, with the transcription factor Sox9 acting as its upstream driver

The normal structure and function of the aortic wall depend on the integrity of the ECM network and the contractile phenotype of VSMCs, which work synergistically to resist shear stress in blood flow. To elucidate matrix and cellular changes in AD, this study included aortic tissue from six patients with AD and six control subjects (Table 1). The Western blot results indicated that in the AD group, the expression levels of SM22α and CNN1, which are markers of the VSMC contractile phenotype, were significantly reduced. Concurrently, the expression levels of the matrix metalloproteinases MMP2 and MMP9 were increased (Fig. 1A-B). The findings indicate a reduction in the VSMC contractile phenotype in the aorta of AD patients, accompanied by degradative remodelling of the ECM. Furthermore, Movat staining revealed a sparse arrangement of medial fibres in AD, significant loss of VSMCs, and a marked accumulation of abnormal proteoglycans in the media region, suggesting that ECM degradation and abnormal matrix deposition coexist in AD (Fig. 1I).

**Fig 1.**
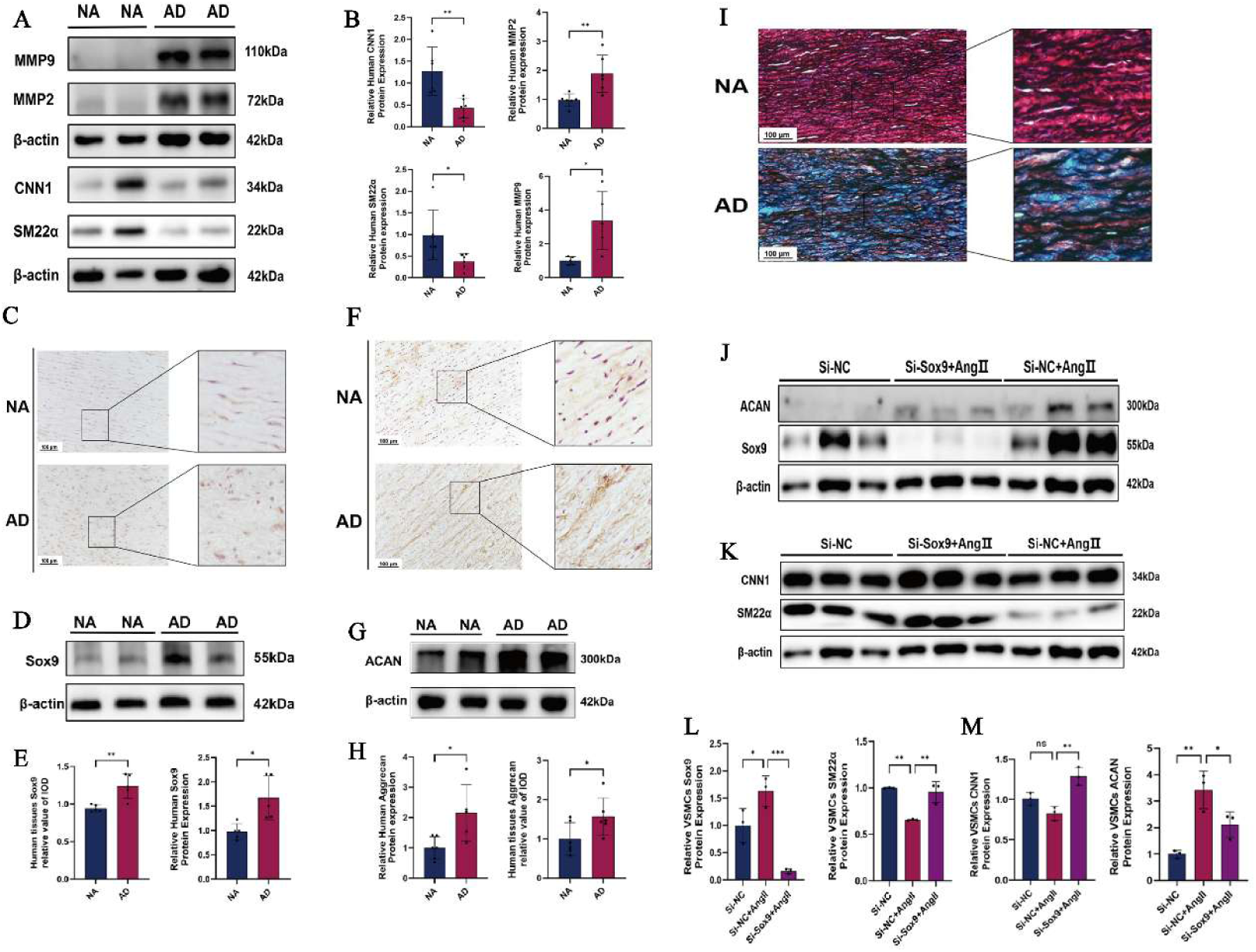
The phenotypic transition of aortic VSMCs in patients with AD is accompanied by proteoglycan accumulation, with the transcription factor Sox9 acting as its upstream driver. A,B, Representative image results from Western Blot showing protein levels of CNN1, MMP2, SM22α and MMP9 in thoracic aortic tissue from healthy controls and AD patients (n=6), with β-actin serving as a super sampling control. C, Representative images of Sox9 protein immunohistochemistry staining from the control group (n=5) and patients with AD (n=5). D, Representative image results from Western Blot showing protein levels of Sox9 protein levels from healthy controls and patients with AD (n=5). E, A semi-quantitative analysis was conducted on the immunohistochemistry and SOX9 Western blot results. F, Representative images of ACAN protein immunohistochemistry staining from the control group (n=5) and patients with AD (n=5). G, Representative western blot images showing ACAN protein levels in thoracic aortic tissue from healthy controls and patients with AD (n=5). H, A semi-quantitative analysis was conducted on the immunohistochemistry and ACAN Western blot results. I, Elastic fibres (black/brown), spindle-shaped vascular smooth muscle cells (red), and intralaminar collagen (yellow) and proteoglycans (blue). J-K, Representative Western blot images showing protein levels of Sox9, ACAN, SM22α and CNN1 in AD tissue, with β-actin used as an internal control. L-M, The statistical outcomes of the investigation into protein expression levels by Western blot are presented (n=3). Statistical analysis was performed using ImageJ and Prism 8.0; all data are expressed as mean ± SEM.

As a pivotal upstream transcription factor for ACAN, Sox9 has been shown to regulate the conversion of VSMCs into chondrogenic/osteogenic cells in chondrocytes and in vascular calcification lesions [28, 29]. However, further research is required to elucidate its expression pattern in AD and its regulation of ACAN. The present study has confirmed, via IHC and Western blot, a significant elevation in Sox9 expression in the aorta of AD subjects (Fig. 1C-E). Furthermore, IHC analysis has revealed that this protein localizes to the nuclei of VSMCs in the media. Concurrently, ACAN expression was significantly upregulated in AD aortas (Fig. 1F-H); IHC staining further confirmed that ACAN was deposited in the medial ECM, consistent with Movat staining results. In view of the histological co-upregulation of Sox9 and ACAN, a further in vitro simulation of the AD microenvironment was conducted by stimulating HAVSMCs with Ang II and knocking down Sox9 using siRNA to validate its regulatory role. The results demonstrated that Sox9 knockdown substantially reduced Ang II-induced ACAN protein accumulation and significantly restored CNN1 and SM22α expression levels. This finding indicates that Sox9 suppression not only diminishes ACAN deposition but also counteracts Ang II-induced VSMC phenotypic transformation (Fig. 1J-M).

### ACAN has been demonstrated to mediate Ang II-induced mitochondrial damage and apoptosis in VSMCs

Abnormal deposition of the protein ACAN in the aorta of patients with AD has been shown to alter the extracellular microenvironment surrounding VSMCs significantly. Recent studies also suggest potential cross-regulation between ECM components and intracellular mitochondrial function. Therefore, the present study investigated whether ACAN is involved in Ang II-induced disruption of mitochondrial homeostasis in VSMCs.

Under normal physiological conditions, the dynamic equilibrium of mitochondrial fission and fusion is maintained. In a previous study, VSMC mitochondria in the pathological state of AD exhibited excessive fission, characterized by up-regulation of fission-related proteins and down-regulation of fusion-related proteins (Mfn1 and OPA1). To elucidate the function of ACAN in this process, ACAN was knocked down in Ang II-stimulated HAVSMCs, and the expression of proteins associated with mitochondrial dynamics was analyzed. The results demonstrated that knocking down ACAN successfully countered Ang II-induced ECM remodelling (Fig. 2A-B), mitochondrial fragmentation (Fig 2C-D), and apoptosis (Fig. 2E-F), while also enhancing mitochondrial membrane potential (Fig 2H-I) and overall mitochondrial health morphology (Fig. 2G). This finding suggests a potential role for ACAN in regulating mitochondrial dynamics.

**Fig 2.**
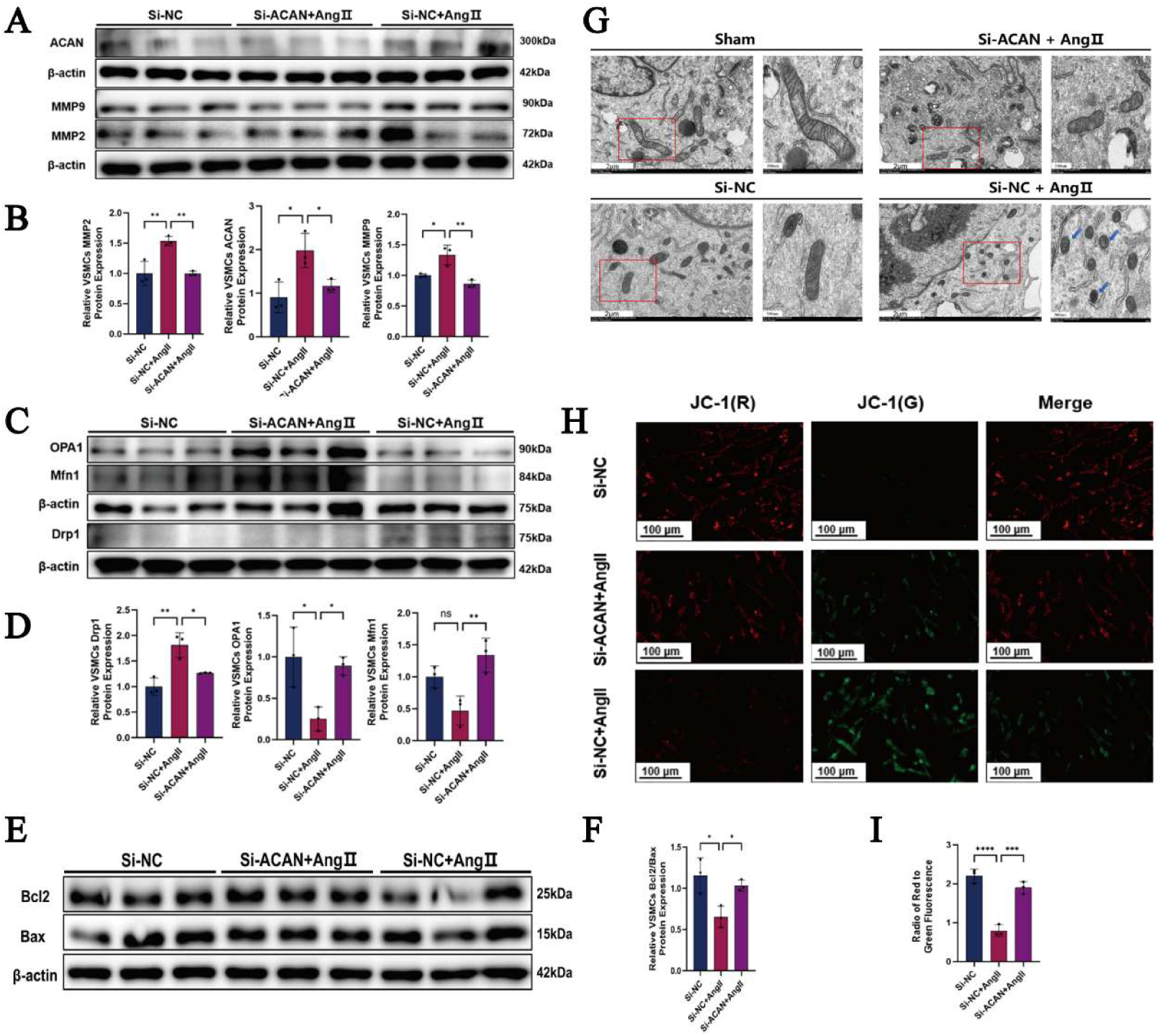
ACAN has been shown to have the capacity to act as a mediator in the process of Ang II-induced mitochondrial damage and apoptosis in VSMCs A, Representative Western blot images illustrating the protein levels of MMP2, ACAN and MMP9 (n=3). **B,** Statistical results of Western blot expression levels (n=3). **C,** Representative Western blot images showing the protein levels of Drp1, OPA1 and Mfn1 (n=3). **D,** Statistical results of Western blot expression levels (n=3). **E,** Representative western blot images illustrating the Bcl2:Bax expression ratios in each group (n=3). **F,** Statistical results of western blot expression levels (n=3). **G,** The mitochondrial morphology of HAVSMCs from each group was examined using a transmission electron microscopy analysis. **H,** JC-1 staining in different groups (n=3). **I,** semi-quantitative fluorescence analysis results (n=3). Statistical analysis of the results was performed using ImageJ software and Prism 8.0. All data are expressed as mean ± SEM.

### In vivo knockout of ACAN has been demonstrated to suppress the onset of AD, a mechanism that is related to the regulation of the Hippo/ Yap1 signalling pathway

In order to extend and validate the findings on ACAN-induced mitochondrial damage in vitro to an in vivo pathological model, an AD mouse model was established using a combination of BAPN and Ang II (Fig. 3A). Furthermore, the expression of ACAN in aortic VSMCs was reduced through the administration of AAV-ACAN via tail vein injection (Fig. 3C-E). At the phenotypic level, mice in the AD model group exhibited marked aortic dilation, with some developing intramural hematomas (Fig. 3B). The incidence of AD reached 66.67%, and deaths began to occur as early as day 21. In contrast, the incidence of AD in the ACAN-knockdown group was significantly reduced to 38.10%, and the first death was delayed to day 24, indicating that ACAN inhibition effectively reduces the risk of AD and improves survival (Fig. 3F-G).

**Fig 3.**
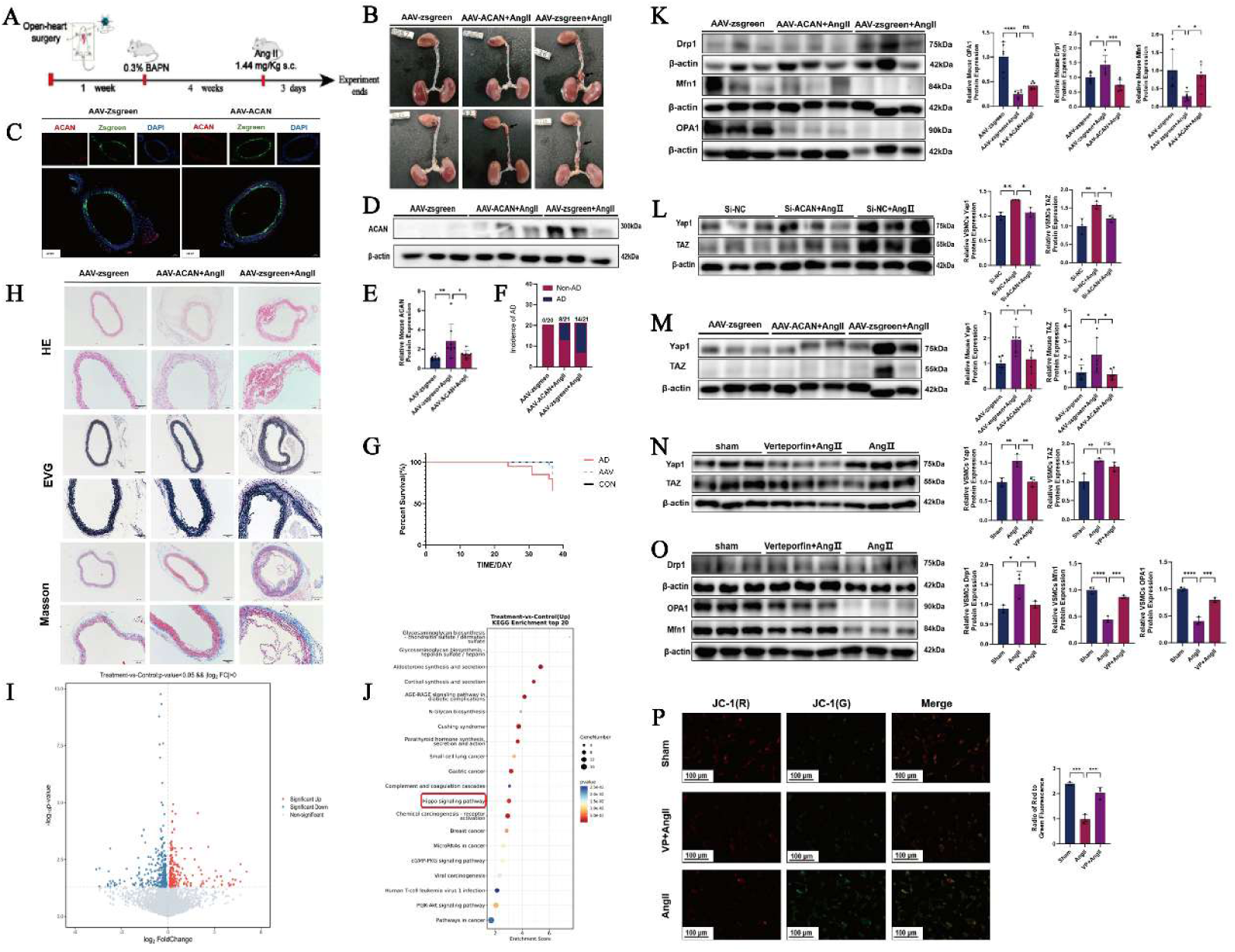
The suppression of the onset of AD has been demonstrated to be a consequence of in vivo knockout of ACAN, a phenomenon that is related to the regulation of the Hippo/YAP1 signalling pathway A, The following flowchart delineates the experimental procedure for establishing a mouse AD model. **B,** The following represents a representative anatomical diagram of a mouse aorta. **C,** Representative fluorescence microscopy images of ACAN (red), zsgreen (green), and DAPI (blue) in the aortas of AAV-zsgreen and AAV-ACAN mice are presented in figure 1. **D,E,** Western blot analysis of ACAN expression in the aortas of the three groups of mice treated with AAV-zsgreen (n=8 each group). **F,** The incidence of AD in the three groups of mice. **G.** The following figure illustrates the survival curves for mice in the various groups. **H,** HE, EVG, and Masson staining. **K,** Representative Western blot images and statistical results for Drp1, OPA1, and Mfn1 protein expression in mouse VSMCs across the groups (n=6). **I,** Volcano plot. **J,** KEGG pathway analysis. **L,** Representative Western blot images and statistical results for Yap1 and TAZ protein expression in VSMCs across the groups (n=3). **M,** The following images illustrate representative western blots and statistical results for Yap1 and TAZ protein expression in mouse aortas across different groups (n=6). **N,** The following images depict representative western blots, alongside the statistical results for Yap1 and TAZ protein expression in VSMCs across various groups (n=3). **O,** Shows representative Western blot images and statistical results for the expression levels of Drp1, OPA1, and Mfn1 proteins in VSMCs across different groups (n=3). **P,** Demonstrates JC-1 staining patterns and semi-quantitative fluorescence analysis results in VSMCs across various groups (n=3). Statistical analysis was performed using ImageJ and Prism 8.0; all data are expressed as mean ± SEM.

Subsequent analysis revealed the multifaceted protective effects of ACAN knockdown on the aortic wall structure. It was revealed that the aortic walls in the AD model group exhibited incomplete structures, characterized by structural disruption and the formation of typical dissections. EVG revealed a significant reduction in the number of elastic fibres, accompanied by disorganized arrangement and tearing. Masson staining revealed excessive collagen deposition (Fig. 3H). In contrast, the aortic lumen in the ACAN knockdown group remained intact, with no intramural hematoma or false lumen formation; elastic fibres were arranged continuously and neatly, and collagen deposition was significantly reduced. Taken together, these results indicate that ACAN knockdown effectively alleviates the vascular wall structural damage and pathological remodelling induced by the combination of BAPN and Ang II.

Concurrently, the regulatory function of ACAN in VSMC mitochondrial homeostasis was validated in vivo. The Western blot results indicated that the aortas of mice in the AD model group showed reduced expression of the fusion proteins Mfn1 and OPA1. In contrast, Drp1 expression was elevated. This finding indicates a pathological state of excessive mitochondrial fission. However, these changes were significantly reversed by ACAN knockdown (Fig. 3K).

Following the initial discovery that ACAN regulates mitochondria in both in vivo and in vitro settings, further investigation was undertaken into its downstream molecular mechanisms. To achieve this objective, RNA-seq was performed on Ang II-stimulated ACAN-knockdown and control HAVSMCs. The results demonstrated that Ang II significantly activated the Hippo signalling pathway. In contrast, ACAN knockdown effectively inhibited its activation, suggesting that the Hippo pathway may be a key mediator of ACAN’s regulation of mitochondria (Fig. 3I-J). Given that Yap1 and TAZ are core downstream effectors of the Hippo pathway, we performed Western blot analysis and confirmed that ACAN knockdown significantly suppressed the Ang II-induced upregulation of Yap1 and TAZ proteins in HAVSMCs; consistent results were also observed in aortic tissues from AAV-ACAN mice, establishing the consistency of the ACAN-Hippo/Yap1 regulatory axis both in vivo and in vitro (Fig 3L-M).

To elucidate the causal relationship between Yap1/TAZ activation and mitochondrial damage, HAVSMCs were treated with the Yap1-specific inhibitor verteporfin. The results demonstrated that Verteporfin not only effectively blocked Ang II-induced upregulation of Yap1 and TAZ, but also significantly downregulated the expression of the fission protein Drp1, while restoring the levels of the fusion proteins Mfn1 and OPA1 (Fig. 3N-O). This finding suggests that the inhibition of the Yap1/TAZ pathway can directly correct Ang II-induced mitochondrial dynamic imbalance. The use of a fluorescent probe, JC-1, to detect mitochondrial membrane potential, confirmed that Verteporfin significantly reversed the decrease in membrane potential induced by Ang II (Fig. 3P).

This finding confirms the essential role of the Yap1/TAZ pathway in sustaining mitochondrial health in VSMCs. Overall, the study shows that ACAN promotes mitochondrial fission and dysfunction in VSMCs by activating the Hippo/YAP1 signalling pathway. Mitochondrial balance can be restored effectively by specifically inhibiting ACAN or blocking the Yap1/TAZ pathway.

## Discussion

This study investigated the impact of increased Sox9 expression on the contractile phenotype of VSMCs and the ECM in AD-induced aortic dissection. The study revealed the mechanism by which ACAN influences VSMC mitochondrial dynamics during AD development, providing a new perspective for further elucidating the pathological mechanisms underlying AD and identifying potential new targets for future AD drug development.

Previous studies on aortic dissection generally focused on changes to the elastic fibres or collagen in the aortic walls of patients with AD. However, recent studies have found that abnormal proteoglycan accumulation in AD may also promote its onset and progression. This accumulation is regarded as a marker of degenerative changes in the aortic intima ^28–30^. In this study, we found that ACAN protein expression was significantly higher in aortic tissue from AD patients and in an AD mouse model created using BAPN and Ang II, compared with the control group. Previous studies have corroborated this finding, with some showing that ACAN and VCAN are the primary glycoproteins elevated in AD. Subsequent research by María et al. demonstrated that inhibiting VCAN expression improves aortic dilation in MFS mice; however, the pathological mechanism underlying ACAN’s role in aortic lesions in AD remains unclear. In this study, we used an adenovirus to specifically inhibit ACAN expression in mouse aortic vascular smooth muscle cells. This improved the survival rate in AD mice and alleviated aortic media lesions, including elastic fibre rupture and collagen deposition. This delayed the onset and progression of AD. These findings suggest that ACAN may be a potential therapeutic target for AD.

The polysaccharide branches of proteoglycans have been demonstrated to play a pivotal role in regulating the dynamic equilibrium of the aortic media and the integrity of the extracellular matrix. At normal physiological levels, proteoglycans support the vascular wall by generating osmotic swelling pressure through the binding of water molecules to their glycosaminoglycan chains. This enables the elastic laminae to resist circulatory compression while placing the elastic fibres and radially oriented microfibrillar cell junctions under tension ^31^. Some researchers have hypothesized that excessive proteoglycans in AD may exert excessive osmotic swelling pressure on the vascular wall, thereby promoting tearing of the vascular media ^11^. Further research has established that the substantial buildup of proteoglycans in AD leads to both mechanical damage and the potential for VSMC demise, which may ultimately culminate in cell death ^29^.

Nevertheless, the precise molecular mechanisms underpinning these processes remain to be elucidated. Moreover, a study was conducted to compare the levels of ACAN in the blood of individuals diagnosed with AD and those who were deemed healthy. The results of this study indicated that individuals diagnosed with AD exhibited elevated levels of ACAN in their blood, suggesting that ACAN could serve as a potential early diagnostic biomarker for AD ^28^.

The validation of clinical samples revealed that patients with AD exhibit reduced expression of contractile phenotype markers in aortic media vascular smooth muscle cells, while matrix metalloproteinases are overexpressed. This indicates a shift of smooth muscle cells from a contractile phenotype to an osteogenic phenotype, accompanied by ECM remodelling. Sox9 is a transcription factor that regulates ACAN expression in cartilage tissue; it is also upregulated in vascular tissues associated with diseases such as atherosclerosis and transplant-associated arteriosclerosis. Current research suggests that Sox9 plays a crucial role in vascular calcification and in the transition of VSMCs toward osteogenic and chondrogenic phenotypes ^32,33^. A recent study reported that Sox9 promotes VSMC phenotypic regulation by modulating collagen expression and reducing VSMC contractility, thereby modulating ECM stiffness ^34^. The present study also found that inhibiting Sox9 expression helps alleviate Ang II-induced VSMC phenotypic transition and elevated matrix metalloproteinase levels, while simultaneously reducing abnormal ACAN expression. This finding suggests that the aberrant upregulation of Sox9 in AD may result in the loss of the contractile phenotype in VSMCs and ECM remodelling. Furthermore, it may contribute to the abnormal accumulation of proteoglycans in AD. A further finding of this study is the influence of ACAN on intracellular mitochondrial dynamics in VSMCs. In recent years, researchers have observed that alterations in the ECM may affect mitochondrial dynamic homeostasis. Mitochondrial homeostasis is imperative for maintaining the normal physiological function of VSMCs. It has been established that normal mitochondria alter their morphology, size, and location through continuous fusion and fission to adapt to changes in the external environment and maintain the cell’s normal energy supply ^35^. Mitochondrial fusion is defined as the process by which two mitochondria sequentially fuse via their outer and inner membranes to reorganize into a single tubular network. This process enhances the overall stability and damage resistance of the mitochondrial network. At the molecular level, the fusion process is synergistically mediated by the Mfn1/2 proteins on the outer mitochondrial membrane and the OPA1 protein on the inner membrane. Mitochondrial fusion has been demonstrated to facilitate the exchange and complementation of mtDNA, membrane phospholipids, and respiratory chain-related proteins within the network. In addition, it has been shown to regulate the metabolic efficiency of the tricarboxylic acid cycle. Conversely, mitochondrial fission fragments the tubular network into smaller organelles, thereby facilitating the elimination of depolarised mitochondria via mitochondrial autophagy ^36^. Mitochondrial fission is primarily activated by the phosphorylation of Drp1, which recruits a series of receptors, including Fis1, Mff, MiD49, and MiD51, to the outer membrane. Finally, Drp1 assembles into a ring-like structure that encircles and compresses the mitochondria, consuming GTP and ultimately producing two separate organelles ^37^. This process not only addresses the cell’s augmented energy requirements but also facilitates the isolation of damaged mitochondria with diminished membrane potential from the broader network, thereby preserving mitochondrial health ^38^. The OPA1 protein is located within the inner mitochondrial membrane, while the protein in question is located on the outer mitochondrial membrane. Mitochondrial fusion has been demonstrated to facilitate the exchange and complementation of mtDNA, membrane phospholipids, and respiratory chain-related proteins within the network.

In addition, it has been shown to regulate the metabolic efficiency of the tricarboxylic acid cycle. Conversely, mitochondrial fission fragments the tubular network into smaller organelles, thereby facilitating the elimination of depolarised mitochondria via mitochondrial autophagy ^36^. Mitochondrial fission is primarily activated by the phosphorylation of Drp1, which recruits a series of receptors, including Fis1, Mff, MiD49, and MiD51, to the outer membrane. Finally, Drp1 assembles into a ring-like structure that encircles and compresses the mitochondria, consuming GTP and ultimately producing two separate organelles ^37^. This process not only addresses the cell’s augmented energy requirements but also facilitates the isolation of damaged mitochondria with diminished membrane potential from the broader network, thereby preserving mitochondrial health ^38^. In physiological contexts, mitochondrial fusion and fission are characterized by a delicate, constantly shifting equilibrium. Disruption of this equilibrium, for instance, by impairment of fusion or fission processes, has been shown to precipitate mitochondrial dysfunction, which can, in turn, lead to the onset of diverse pathological conditions. Mitochondria in VSMCs have been demonstrated to play a critical role in maintaining aortic homeostasis. A growing body of evidence indicates that alterations in mitochondrial dynamics and biogenesis may be substantial pathological features of AD ^25,39^.

In AD model mice established by the combination of BAPN and Ang II, as well as in Ang II-induced VSMC models, Drp1 expression was found to be increased, while Mfn1 and OPA1 expression levels were reduced. This indicates that mitochondrial homeostasis is disrupted and excessive mitochondrial fission occurs during the development of AD. Inhibition of ACAN expression in mouse aortas and VSMCs has been shown to enhance mitochondrial function, as evidenced by subsequent RNA-seq analysis, which revealed that ACAN knockdown alters Hippo pathway expression. The Hippo-YAP/TAZ signalling pathway has been identified in a variety of human tissues, including the nucleus, cytoplasm, and muscle. Yap/TAZ has been identified as a sensor and mediator of mechanical cues arising from extracellular matrix stiffness, cell geometry, cell density, and actin cytoskeletal status. Among these, Yap1, a pivotal factor in mechanotransduction, has been shown to regulate cell survival and proliferation by binding to DNA-binding transcription factors, thereby inducing gene expression ^40^. ACAN, a proteoglycan, has been hypothesized to influence the expression of the transcription factor Yap by modulating the mechanical properties of the ECM. A recent study revealed a link between the Yap1/TAZ pathway and mitochondrial fission and fusion homeostasis; targeting the Yap1/TAZ axis can alter mitochondrial homeostasis ^41^. Consequently, ACAN may influence mitochondrial dynamic homeostasis by regulating the Yap1/TAZ pathway. To validate our hypothesis, we demonstrated the effect of ACAN on the Yap1/TAZ pathway in both in vivo and in vitro settings. After the inhibition of Yap1 expression using the Yap1 inhibitor verteporfin, an increase in mitochondrial membrane potential was observed in VSMCs under AD conditions, indicating improved mitochondrial function.

In summary, abnormally elevated Sox9 levels in the aortic wall promoted ACAN accumulation, which in turn induced excessive mitochondrial fission in VSMCs by activating the Yap1/TAZ pathway, thereby contributing to the onset and progression of AD. These findings provide new insights into the mechanisms underlying the onset and progression of AD.

## Conclusion

In view of the hypothesis that proteoglycans may play a pivotal role in the pathogenesis of AD, the aggregating proteoglycan ACAN was selected as the research subject. The close association between ACAN and AD further confirmed this selection.

1. ACAN protein levels were found to be increased in the aortic tissues of AD patients and in AD mice induced by BAPN combined with Ang II.
2. Inhibition of Sox9 has been demonstrated to attenuate VSMC phenotypic transition and ECM remodelling.
3. Inhibiting the abnormally elevated expression of ACAN has been demonstrated to help mitigate the development of AD in mice and improve survival rates.
4. Elevated ACAN expression has been shown to induce excessive mitochondrial fission via the Yap1/TAZ pathway, leading to vascular smooth muscle dysfunction and resulting in AD.
5. Verteporfin has been demonstrated to mitigate the development of AD by inhibiting the Yap1/TAZ pathway, thereby reducing excessive mitochondrial fission and improving mitochondrial dysfunction.

## Author Contributions

Wenjie Fu, Yanlan Gao: Writing—Original Draft. Yanqing Fang, Linlong Guo, Yalu Huang, Luyao Hong: Data curation. Ziru Li, Bingxing Lu, Laixiang Yan, Bin Hu: Investigation. Chaoyun Wang: Supervision. Lin Zhong: Methodology. Yumei Li and Min Chen: Writing— Review and Editing.

## Acknowledgments

The authors thank the Union Hospital of Fujian Medical University and the Public Technology Service Center of Fujian Medical University for their support of the experiments. The authors thank the patients and medical staff for their support of this study.

## Data availability statement

The data that support the findings of this study are available from the corresponding authors upon reasonable request.

